# Real-Time Spatial Registration for 3D Human Atlas

**DOI:** 10.1101/2022.09.27.509335

**Authors:** Lu Chen, Dejun Teng, Tian Zhu, Jun Kong, Bruce W. Herr, Andreas Bueckle, Katy Borner, Fusheng Wang

## Abstract

The human body is made up of about 37 trillion cells (adults). Each cell has its own unique role and is affected by its neighboring cells and environment. The NIH Human BioMolecular Atlas Program (HuBMAP) aims at developing a 3D atlas of human body consisting of organs, vessels, tissues to singe cells with all 3D spatially registered in a single 3D human atlas using tissues obtained from normal individuals across a wide range of ages. A critical step of building the atlas is to register 3D tissue blocks in real-time to the right location of a human organ, which itself consists of complex 3D substructures. The complexity of the 3D organ model, e.g., 35 meshes for a typical kidney, poses a significant computational challenge for the registration. In this paper, we propose a comprehensive framework TICKET (<u>TI</u>ssue blo<u>CK</u> r<u>E</u>gis<u>T</u>ration) for tissue block registration for 3D human atlas, including (1) 3D mesh pre-processing, (2) spatial queries on intersection relationship and (3) intersection volume computation between organs and tissue blocks. To minimize search space and computation cost, we develop multi-level indexing on both the anatomical structure level and mesh level, and utilize OpenMP for parallel computing. Considering cuboid based shape of the tissue block, we propose an efficient voxelization-based method to estimate the intersection volume. Our experiments demonstrate that the proposed framework is practical and efficient. TICKET is now integrated into the HuBMAP CCF registration portal [1].

## 1 INTRODUCTION

With the significant development of tissue imaging technologies[2] [3] [4] [5], enormous amounts of information about our human body are now produced at both cellular and subcellar levels. The Human BioMolecular Atlas Program (HuBMAP) [6] of National Institutes of Health is a national initiative to develop an open, global framework for comprehensively mapping of the human body at cellular resolution, in particular, generating foundational 3D tissue atlases of human body, to better understand how the healthy human body works and what changes during aging or disease. A Human Body Common Coordinate Framework [7], encompassing 3D organization of whole organs and thousands of anatomical structures, are being built by HuBMAP, where the organs and the anatomical structures are represented in mesh based 3D models [8].

By collecting 3D tissue blocks from human body, a vast amount of spatial information (cells and biomarkers) will be derived from various tissue imaging technologies on the tissues (2D and 3D). An essential component of HuBMAP is tissue block registration [9] (Figure 1), a critical step to enable organ experts to register tissue data to the atlas. It will also help researchers to explore tissue data in a spatially and semantically explicit manner. In order to explore and analyze the inner structures inside the tissue block, 3D computational geometry and spatial data management plays a major role for exploring spatial relationships, e.g. anatomical structures that are colliding with the tissue block, as well as quantitative measurements, e.g. volume of each anatomical structure inside the tissue block. However, there are several challenges we need to address:

**Figure 1:**
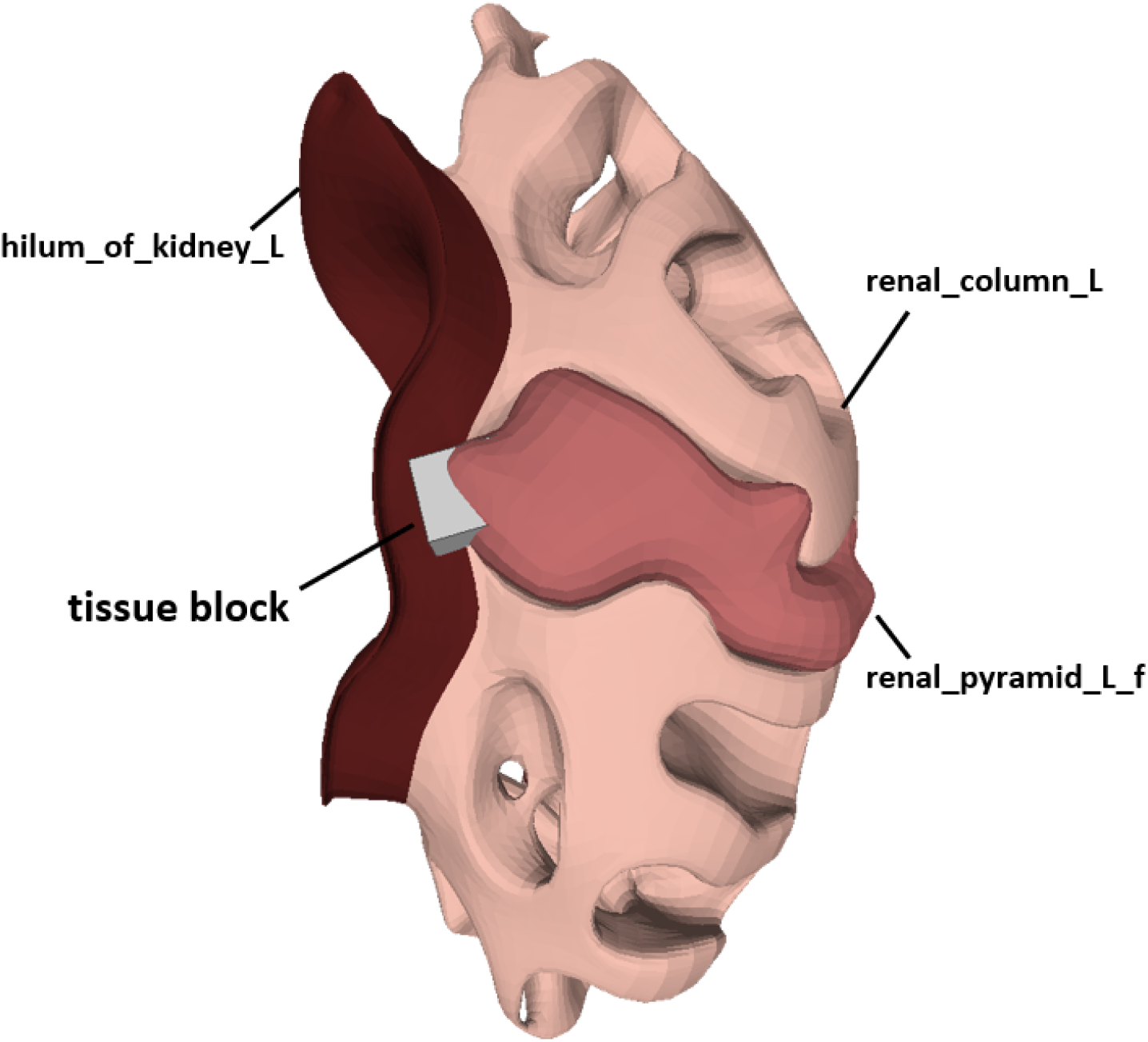
Tissue block registration. The hilum of kidney, renal column and renal pyramid are anatomical structures inside the kidney, which may collide with the tissue block.

### Complex 3D Structures

3D anatomical structures have diverse shapes. Some are in regular shapes while others may have complex structures such as vasculatures and skins as Figure 2 shows. Although minimal bounding boxes (MBBs) is one of practical approaches to simplify complicated models, MBBs is not effective to represent such complex 3D objects for spatial queries. A new indexing method for primitives of 3D models is necessary to reduce the query complexity. Besides, the hierarchical organization of human body demands a multi-level indexing approach.

**Figure 2:**
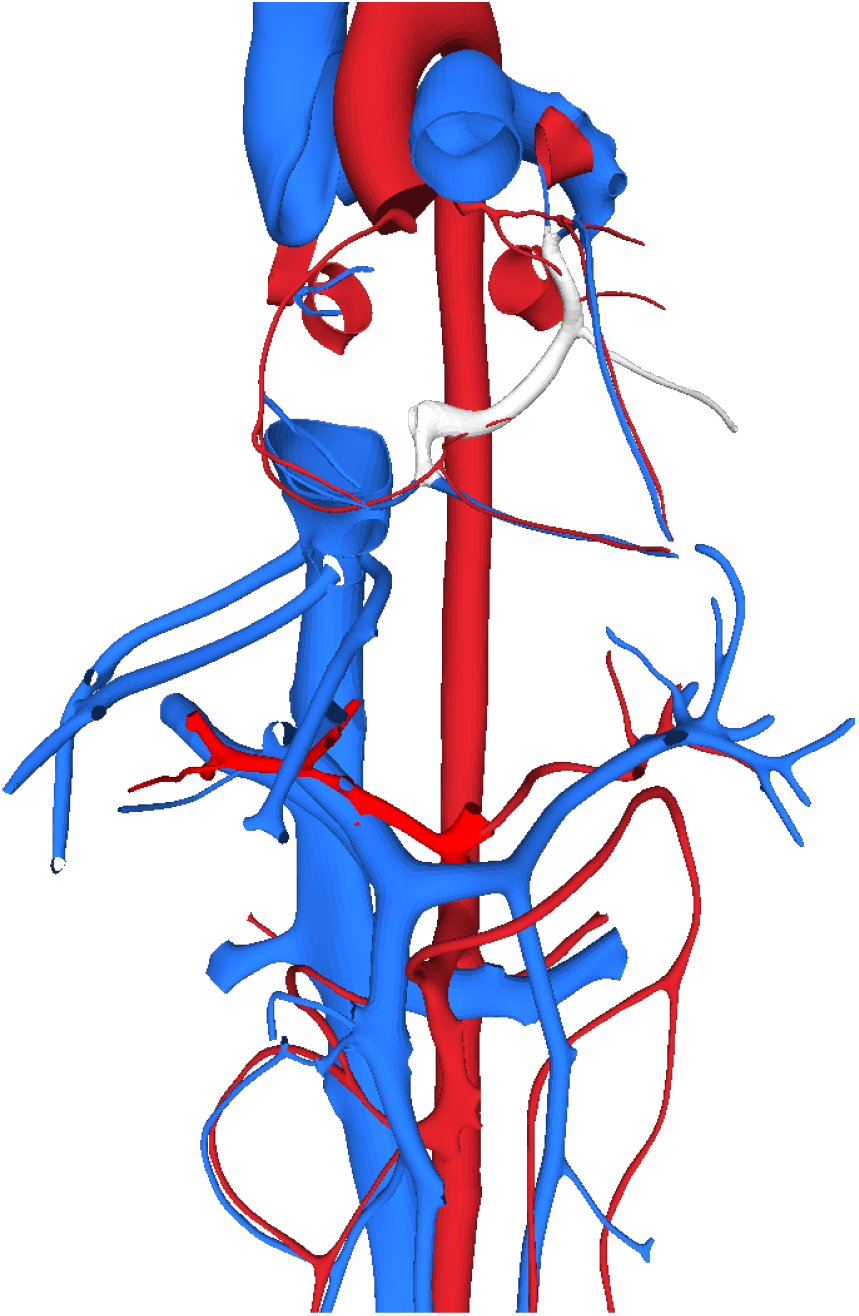
3D models of vasculatures

### Varying Mesh Quality of 3D models

3D models of organs come from different sources such as Allen Brain Atlas [10], the Human Tumor Atlas [11], HuBMAP [6], the Kidney Precision Medicine Project [12] and so on. The mesh quality varies due to different purposes of the projects and varying requirements. For example, visualization oriented 3D organ models are targeted to visualization only, and may have limitations for spatial processing and queries, for instance, existence of non-manifold edges [13] and holes, even though smoothness and other post-processing techniques will make the models “look” smooth and perfect. For scientific computing purpose, 3D models for anatomic structures must be spatially compliant to ensure accurate calculation of spatial relationships and measurements. However, it is inevitable to lose vertices or edges due to the occlusion, reflectance or raw data preprocessing, rendering holes in the surface and non-manifold meshes. For example, the mesh in Figure 3a is divided into two parts, i.e., the renal pyramid and the renal papilla, according to biological functionalities and structures. The holes are automatically generated since the surface is not closed any more due to the division of the mesh in Figure 3b. Therefore, a hole filling algorithm will be applied to the 3D organ models as Figure 3 shows in advance to guarantee the precision of tissue block registration.

**Figure 3:**
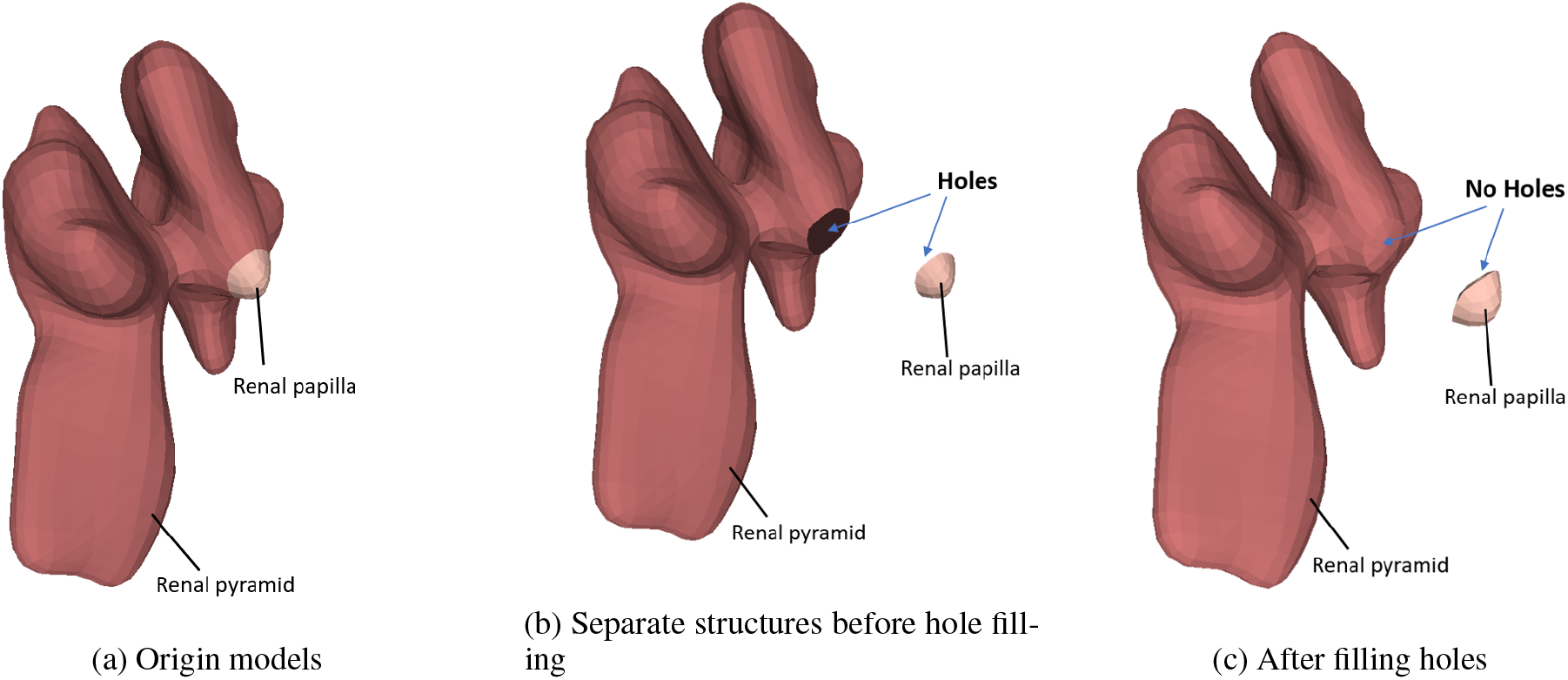
Example of mesh-preprocessing: hole filling

### High Computation Complexity

3D spatial queries involve computationally intensive geometric operations for quantitative measurements and identifications of spatial relationships. While a traditional minimal bounding box (MBB) approach can filter out the candidates dissatisfying the constraints of spatial queries in the filter stage, the refinement stage still cost too much computation when examining the candidates with respect to their exact geometry to generate the exact solutions [14] [15]. Due to the complex structures of 3D objects, the refinement stage could dominate the spatial query cost. Moreover, queries which require quantitative computations such as intersection volume of objects, have much higher complexity than normal query types.

The above challenges demand a practical and efficient framework that can normalize meshes, exploit indexing techniques suitable for complicated objects, reduce computational complexity, and mitigate potential I/O cost to tackle the task of 3D tissue block registration.

Our main contributions are summarized as follows:

- We propose a comprehensive framework TICKET for tissue block registration including three steps: (1) Mesh pre-processing such as remeshing non-manifold meshes and filling the holes to produce watertight meshes, (2) identify the anatomical structures that are collided with the tissue block, (3) compute the volume of each anatomical structure inside the tissue block, if intersected;
- We introduce multi-level spatial indexing at the anatomical structure level and mesh level, which improve query performance compared with the traditional filter-refine paradigm [16]. We take an in-memory approach for data management and query to mitigate the I/O cost and reduce latency. Besides, we provide a spatial query *http* service for tissue block registration, which can be invoked on-demand when users upload the tissue blocks.
- We develop an efficient voxelization-based method to estimate the intersection volume between the tissue block and the organ considering the regular cuboid shape of the tissue block. We utilize parallel programming to accelerate the volume computation.

The rest of the paper is organized as follows. We first introduce the background in Section 2. The framework of tissue block registration is demonstrated in Section 3. Next, we introduce multi-level spatial indexing in Section 4. Section 5 presents the voxelization-based method to estimate the intersection volume as well as the parallel computing scheme to accelerate the volume computation. The performance of the framework is evaluated in Section 6 followed by conclusion.

## 2 BACKGROUND

### 2.1 Mesh Pre-processing

Mesh pre-processing is a fundamental step of 3D data acquisition. Poor quality meshes may cause errors in spatial queries or lead inaccuracy intersection volume computation. There are some typical mesh pre-processing techniques such as mesh self-intersection checking, remeshing and hole filling. Self-intersection checking is intuitive and easy to implement while hole filling is much more subjective and semantic.

Due to the differences of data representation, current hole filling algorithms [17] can be divided into two categories: point cloud-based methods and mesh-based methods. Point cloud-based methods were developed in earlier years, while mesh-based methods gain more attention and play an important role with the rapid development of computer graphics. Typical workflow of hole filling algorithms can be summarized in two steps: hole boundaries should be first detected and then holes are filled by generating new meshes. In particular, mesh-based hole filling algorithms are divided into two categories: volume-based algorithms and surface-based algorithms according to the data used in the phase of filling holes. Normally, the simple triangulation algorithms work well to triangulate the polygonal hole regions, especially for the flat and disk-like holes without intense changes to the hole topology and geometry. For large holes that are too complex to fill using triangulation, various hole filling algorithms are proposed [18] [19] [20] [21]. In this paper, we adopt the hole filling algorithm [20] to fill all the problematic holes.

### 2.2 Spatial indexing

Each human organ consists of a set of anatomic structures (AS). For example, a kidney has renal pyramids, renal papilla, outer cortex, a total of 35 ASs. A brain contains more than 200 anatomical structures. The organ consisting of multiple ASs is represented by a 3D model, typically using a mesh based representation, using formats such as Graphics Language Transmission Format (e.g., GLB) [22] or Object File Format (OFF) [23]. A typical kidney may be represented by 35 mesh surfaces. To register a human tissue block to an organ, proper indexing is needed for such complex 3D models to minimize the search space and geometric computations [24]. The key principle of the spatial indexing is to divide the search space into regions. There are two common types of space decomposition:

- **Regular decomposition**, also known as space driven index, divides the space in a regular manner independent of the distribution of the objects (i.e., indirectly related to the objects in the space). Objects are mapped to the cells according to some geometric criteria. The most common methods are Quad Tree [25] and cell-based Hashing [26].
- **Object Directed Decomposition** is also known as data driven index. The division of the index space is determined directly by objects. The space is divided by one of or a combination of the following properties: coordinates of data points, the extents, the bounding box or the spatial objects. The most common examples are K-D tree [27], BSP-tree [28] and R-tree [29]. Although K-D tree and BSP-tree are easier to implement, the deletion and insertion of them are non-trivial while R-tree is designed for dynamic changing data and support deletion and insertion.

Traditional filter-refine paradigm with single spatial index built on MBBs of objects is not suitable for complex objects because the geometric computation during the refinement stage will dominate the spatial query cost. Therefore, building index on primitives (vertices, edges and faces of meshes) can greatly reduce the complexity.

## 3 OVERVIEW

### 3.1 Framework of the Tissue Block Registration

We aim to develop a comprehensive framework for tissue block registration, which supports mesh pre-processing, spatial queries on intersection relationship and intersection volume computation. We present the overall framework for the tissue block registration in Figure 4. Initial 3D mesh data is stored in disks in object file format (OFF). We first check whether the meshes are 2-manifold or not. If not, the medical illustrators of HuBMAP will remesh the non-manifold meshes manually. The meshes with holes are then pre-processed by the hole filling algorithm [20]. After the hole filling, the generated patches can be refined and faired [30]. Each individual mesh of anatomical structures will be loaded and stored in memory. We also create a hash map to store the meshes, where the key is the organ name and the value is a list of the meshes of the anatomical structures of the organ for an easy search. Next, an anatomical structure level index will be generated according to the MBBs of the anatomical structures for each organ, and a mesh level index is built for the mesh of every anatomical structure. In addition, different from traditional intersection volume computation based on Boolean Operations [31] on meshes, we propose a voxelization-based method to estimate the intersection volume between the tissue block and the anatomical structure. To accelerate the intersection volume computation, we exploit parallel programming framework OpenMP to check whether each voxel of the tissue block is inside the polygon mesh of the anatomical structure in parallel.

**Figure 4:**
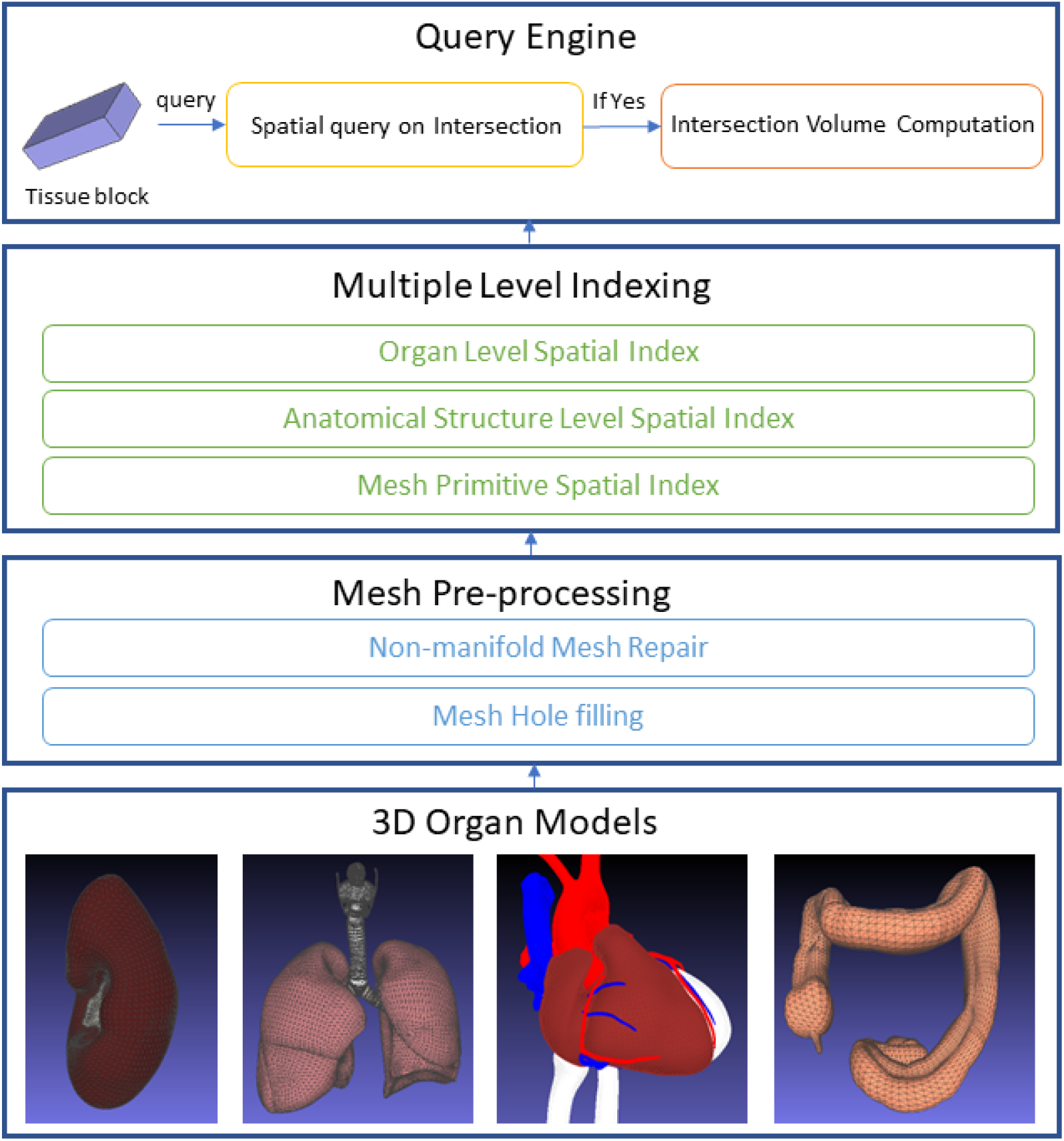
Framework of 3D tissue block registration

### 3.2 On-demand Spatial Query Service

We provide an on-demand spatial query *http* service for the ease of use. Users can send a *http* request with the location information of the tissue block including translation, rotation and scaling parameters. The server will process the request accordingly, with the following procedure (the workflow of the tissue registration is also summarized in Algorithm 1): first transform the rotation angle-based representation of the tissue block to a mesh-based representation. Next, the anatomical structures whose MBBs are not overlapped with the MBBs of the tissue block are filtered out by the anatomical structure level index. The exact intersected anatomical structures with the tissue block are reported by using mesh level spatial index. If intersected, we will compute the intersection volume by proposed voxelization-based intersection volume computation. Otherwise, we will return 0 immediately because they have no intersection.

#### Algorithm 1

Workflow of tissue block registration

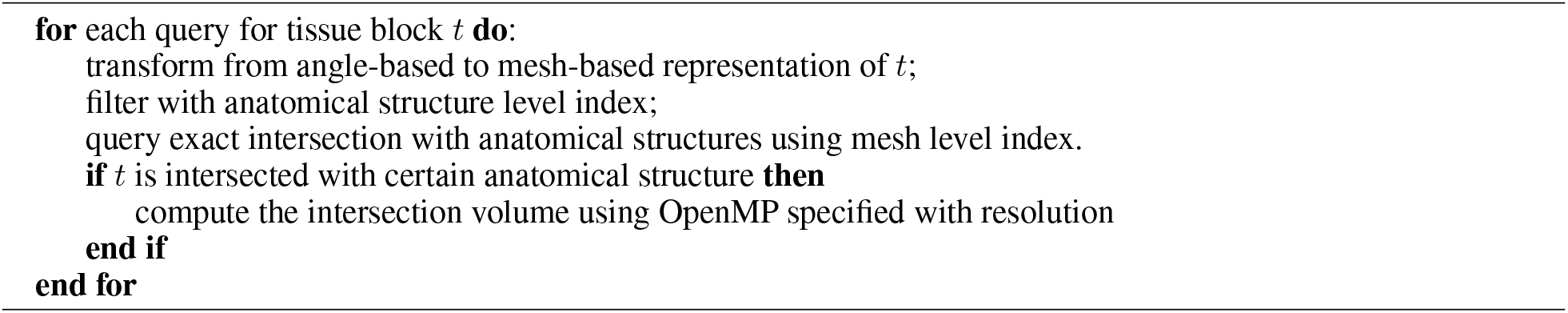

## 4 MULTI-LEVEL SPATIAL INDEXING

### 4.1 Anatomical Structure Level Indexing

Figure 1 demonstrates an organ with internal anatomical structures. Different anatomical structures contain different tissue types and cells, with different functionalities. To fast locate the right positions in the organ for the tissue block registration, we build an anatomical structure level index according to the MBBs of the anatomical structures. An R-tree is constructed on the MBBs of the anatomical structures to achieve the dynamic insertion and deletion of the anatomical structures as Figure 5a presents. If cell models are provided in the future, we can further build spatial index on cells [32] based on cell types and spatial distirbution patterns. Therefore, from macro level to micro level, we build multi-level spatial indexes according to the hierarchical organisation of our human body.

**Figure 5:**
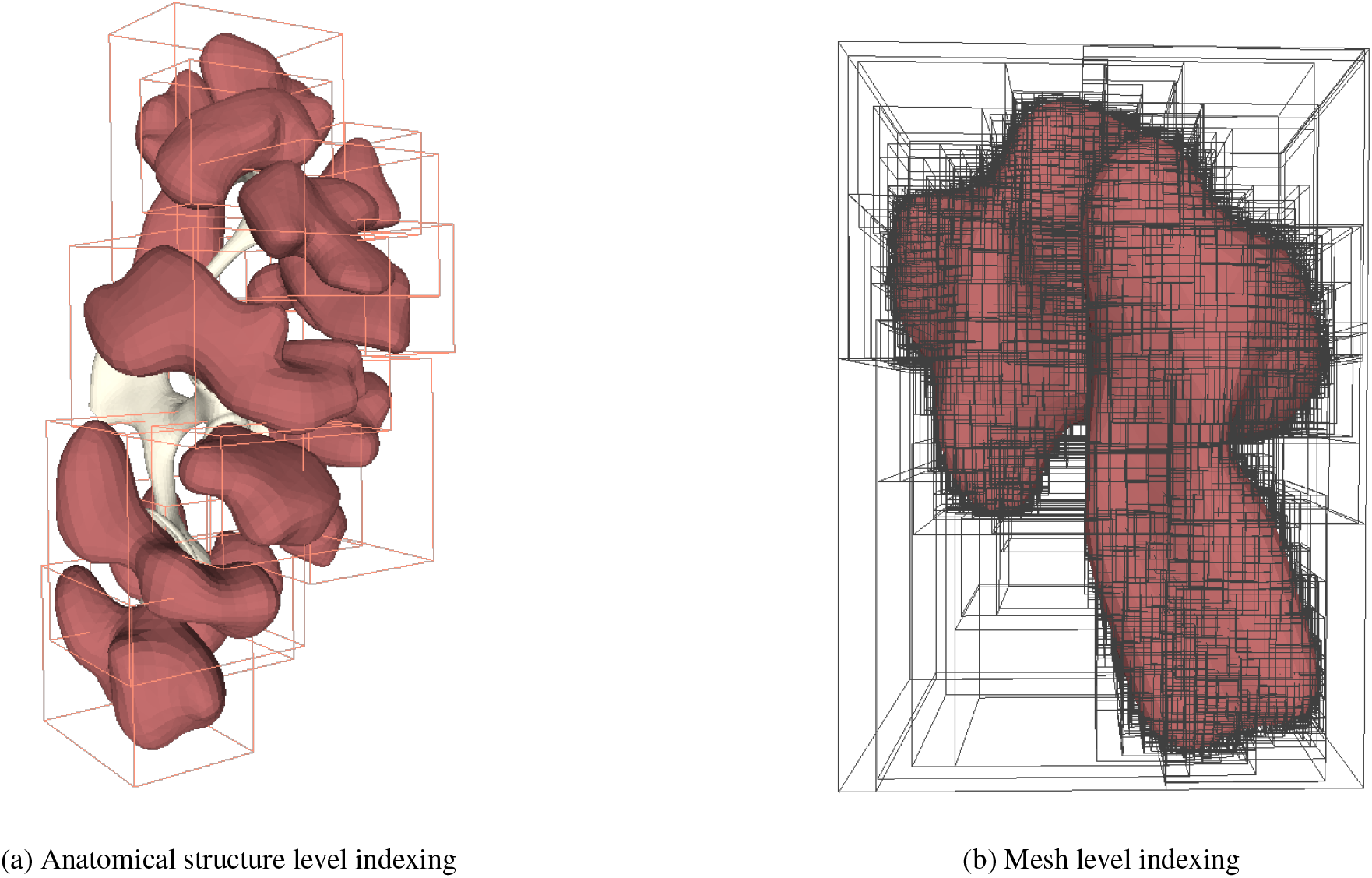
Multi-level spatial indexing

### 4.2 Mesh Level Indexing

For complex structures, traditional filter-refine paradigm suffers from geometric computation in the refinement stage, because the simplified MBBs cannot filter out most unqualified candidates, such as a tissue block outside but near the vessel. Therefore, the mesh-level index built on the primitives (faces) of the complicated structure is necessary as Figure 5b. As the resolution of 3D models becomes higher, a mesh-level spatial index can greatly reduce the complexity of spatial queries from *O*(*N*) to *O*(log *N*) where *N* is the number of primitives. Therefore, we build a BSP-tree, also called AABB-tree on the primitives for every static mesh of the anatomical structures.

## 5 VOXELIZATION-BASED INTERSECTION VOLUME COMPUTATION

The traditional way to compute the intersection volume between two polygon meshes is through Mesh Boolean Operations [31], which consists of two steps: the first step is to compute the mesh intersection of the two polygon meshes. The volume of the intersection mesh is computed next. However, the complexity of Boolean Operations is very high. One of the improvements of Boolean Operations is to use Constructive Solid Geometry (CSG) [33]. Triangle-triangle intersections need to be computed in advance in CSG, which is very time-consuming.

In this paper, we propose an efficient voxelization-based method supporting different resolution of voxelization to estimate the intersection volume. Furthermore, we use the parallel programming framework OpenMP to make the full use of the multi-core processor, which is widely used in modern computers.

### 5.1 Tissue Representation Transformation

Tissue blocks provided by HuBMAP are described with size, rotation, translation, and scaling parameters, which are not the standard mesh representation by vertices, edges and faces. Therefore, we need to recover the mesh from the parametric representation.

Assuming a tissue block is described as:

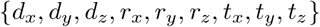

where *d*_*x*_, *d*_*y*_, *d*_*z*_ are the lengths of the tissue block in x,y,z dimensions respectively, *r*_*x*_, *r*_*y*_, *r*_*z*_ are the rotation angles and *t*_*x*_, *t*_*y*_, *t*_*z*_ are the translations. The initial center of the tissue block is the origin and the default rotation order is *xyz* as specified by HuBMAP. Since it is the Euler angle system rather than the fixed angle system that is adopted during rotation by HuBMAP, the rotation order should be adjusted to *zyx* instead. Hence, the rotation matrix with the rotation angles *r*_*x*_, *r*_*y*_, *r*_*z*_ can be written as:

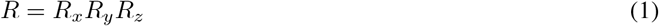

where

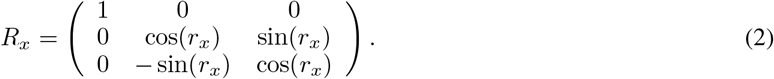

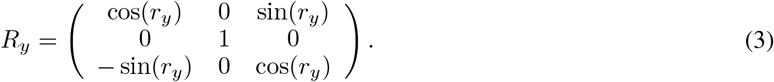

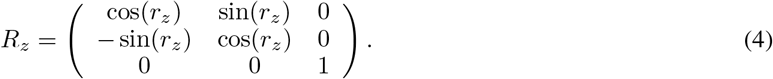

Any point *p* = (*p*_*x*_, *p*_*y*_, *p*_*z*_)^*T*^ inside or on the boundary of the initial tissue block, i.e.,

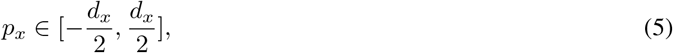

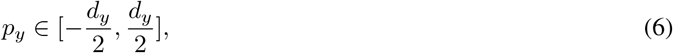

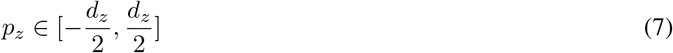

can be transformed by applying the affine transformation: *p*^*′*^ = *Rp* + *T* where *T* = (*t*_*x*_, *t*_*y*_, *t*_*z*_)^*T*^, *R* = *R*_*x*_*R*_*y*_*R*_*z*_. Note that the edges and faces relations, the size of the tissue block are not changed due to the affine invariance. Through the above affine transformation, it becomes fairly simple to construct the standard mesh given the initial coordinates of vertices of the tissue block, the rotation matrix and the translation vector.

### 5.2 Voxelization

A voxel represents a regular grid in 3D space similar to a pixel in a 2D image. Traditionally, voxelization is applied to 3D models in fixed axes directions. Due to regular cuboid shape of the tissue block, we always voxelize the oriented tissue block in its coordinate system along three different diagonal directions. For each voxel, we test whether the center of the voxel lies inside a certain anatomical structure. The estimated intersection volume can be calculated through the volume sum of the voxels whose center are inside the anatomical structure. Via this approach, the complicated volume computation problem is reduced to a point location test, which saves the computational cost greatly.

Point location test is a very classic problem in computational geometry and the current implementation is based on the number of triangles intersected by a ray with the query point as the source. If the number of triangles intersected is odd, the point is inside the polyhedron. Besides, the AABB tree constructed on each mesh can accelerate the point location queries because the ray casting is an intersection problem essentially. The reuse of AABB trees significantly reduces the overhead. To make the full advantage of multi-core processors, we implement the voxelization-based volume computation in parallel using OpenMP. In addition, to avoid memory overflow, we process the voxelization and point location test on the fly rather than store all the voxels in the memory.

Although the complexity of point location query is *O*(log *N*), a higher resolution voxelization requires more computational resources. We define the resolution coefficient *k* as the number of voxels in one unit in one dimension, which indicates a tissue block is divided into *kd*_*x*_ *× kd*_*y*_ *× kd*_*z*_ voxels. We assume that the lowest resolution is *k* = 1 in which the tissue block is divided into *d*_*x*_ *× d*_*y*_ *× d*_*z*_ voxels. Since *d*_*x*_, *d*_*y*_, *d*_*z*_ are integers defined by HuBMAP, we can ignore the “partial” voxel problem. The voxelization-based intersection volume computation algorithm is summarized in Algorithm 2.

#### Algorithm 2

Parallel computing for voxelization-based volume computation

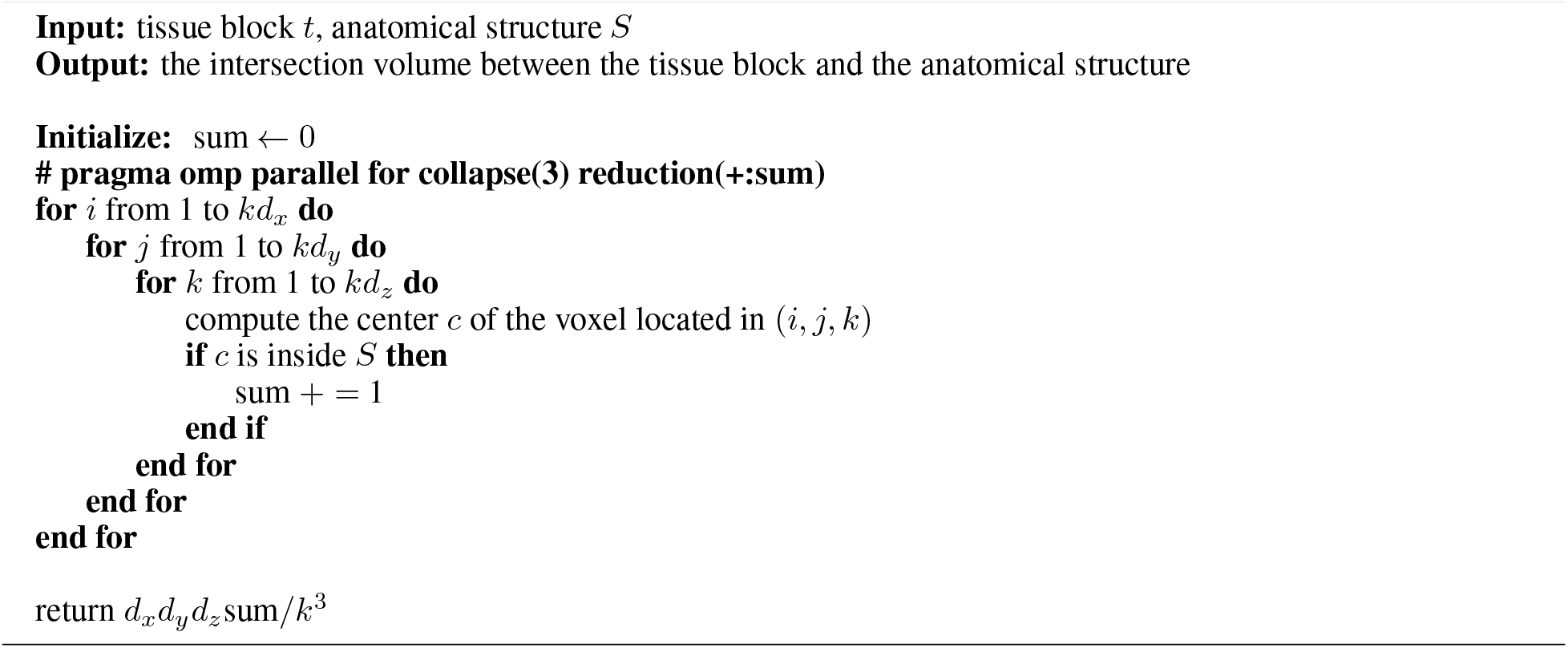

## 6 EXPERIMENTAL RESULTS

All the tests are conducted on a compute node with a 24-core CPU (AMD Ryzen Threadripper 3960 at 3.8GHz). The node comes with 128GB memory(DDR4 3200) and 2TB SSD (NVMe M.2 PCI-Express 3.0). The OS is Ubuntu 20.04.4 with 5.4.0-124 kernel.

We conduct these tests over the official HuBMAP 3D reference organ models [8] and tissue block dataset [34], which contains 68 tissue blocks registered in kidneys.

First, we compare the loading time of the models and the time of building multi-level spatial indexes for five organ models, namely Kidney, Allen Brain, Lung, Heart and Small Intestine. From Figure 6, we can see that large models such as Allen Brain which contains 282 anatomical structures with total of 324,833 vertices and 656,268 faces, cost more time to load and build spatial indexes while small models such as Small Intestine which contains 9 anatomical structures with total of 16998 vertices and 33524 faces, cost less time to load and build spatial indexes. Besides, building spatial indexes costs only half of the time of loading 3D models to the memory.

**Figure 6:**
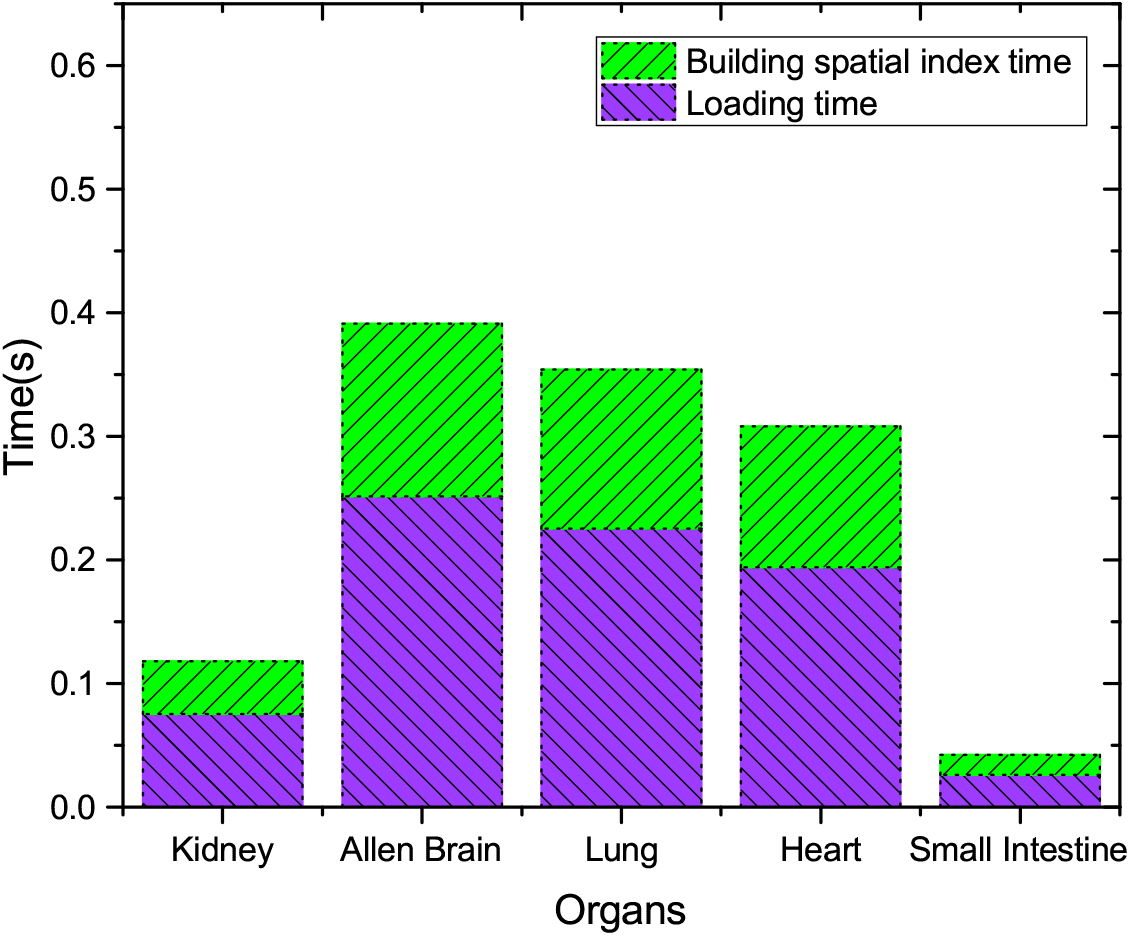
Loading time and building indexes time of different organs

Next, we evaluate four different methods (brute force, only building index on MBBs of anatomical structures, only building spatial index on primitives for each mesh, and building multi-level spatial index both on MBBs of anatomical structures and primitives) of the spatial query on intersection. Figure 7 shows the total query time of all 68 tissue blocks of the spatial query on intersection under four different methods. It is obvious that the spatial index on MBBs of ASs, which follows the traditional filter-refine paradigm, can greatly reduce the query time compared with the brute force method. Another important fact is that the spatial index built on primitives can further reduce the query time. We also test the spatial intersection query using PostGIS, which is extremely slow. Therefore, the multi-level spatial indexing can significantly improve the performance compared with all other methods.

**Figure 7:**
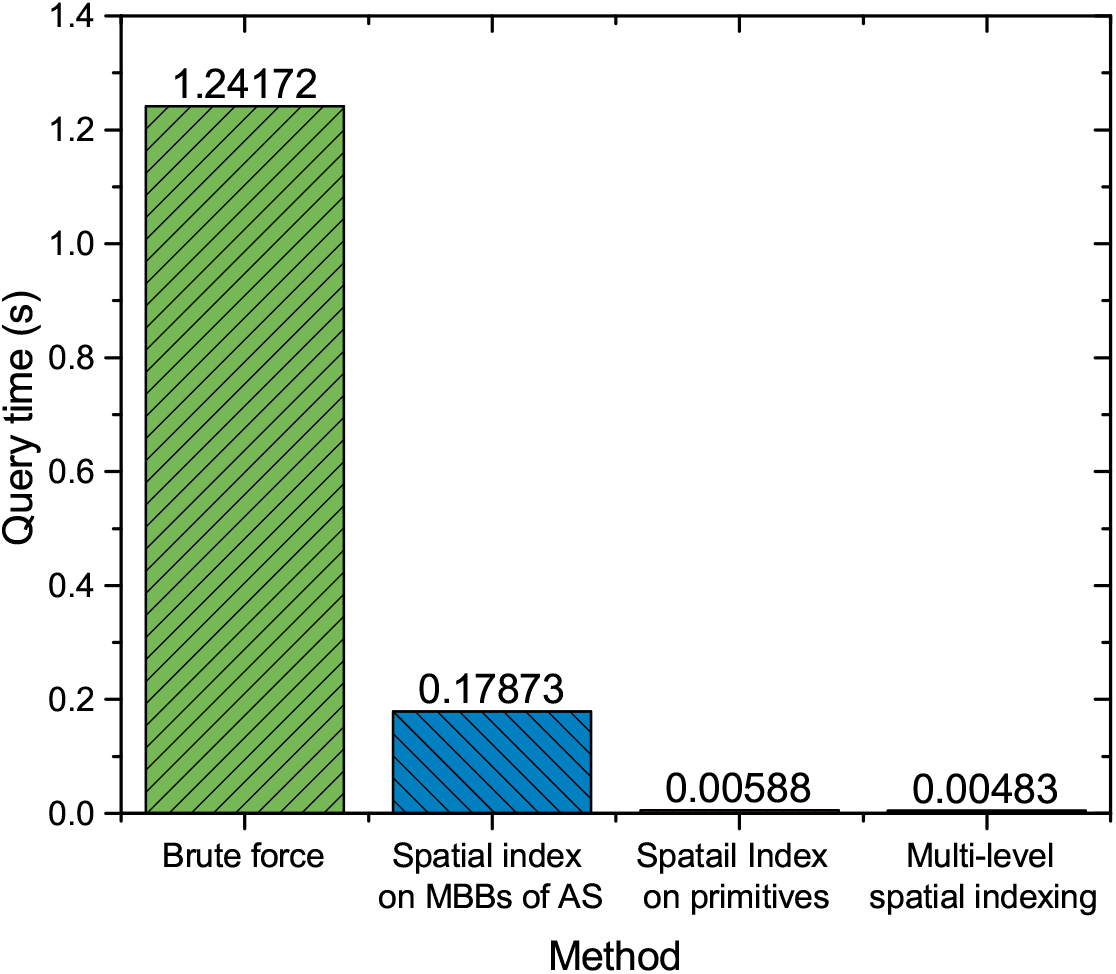
Comparisons of different indexing methods

Moreover, we investigate the impact of parallel computing using OpenMP on voxelization-based intersection volume computation method. From Figure 8 we can conclude that the overhead of cache and synchronization of parallel computing results in no improvement in terms of the query performance when the resolution is low. The number of parallel units increases when the resolution increases. Therefore, the query can make full advantage of multi-core processor, which greatly reduce the query time compared with serial computation.

**Figure 8:**
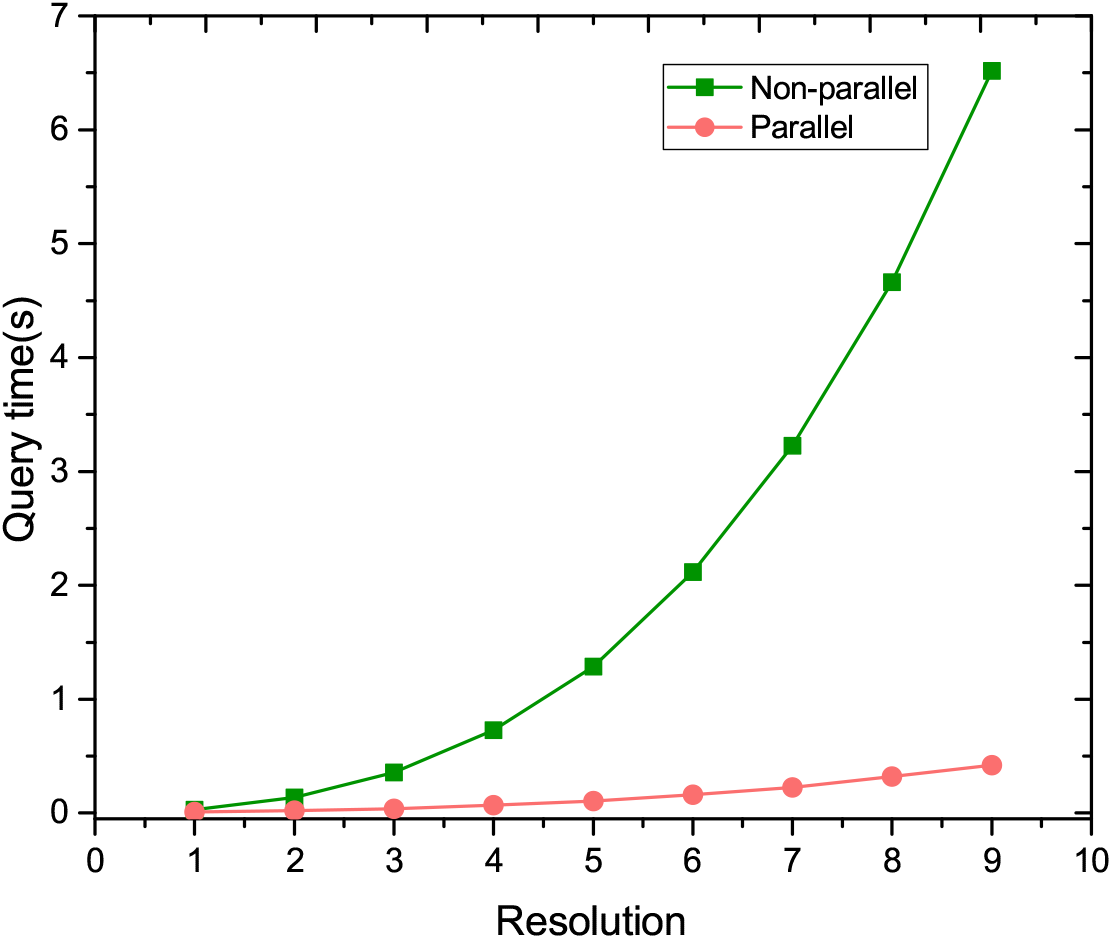
Comparisons of query time between parallel and non-parallel

Finally, we demonstrate the accuracy of the proposed voxelization-based intersection volume computation method. We use the relative error in most cases where the interaction volume exceeds a threshold. Meanwhile, we use the absolute error instead when the intersection volume is around 0. As Figure 9 shows, the error rate decreases as the resolution increases. The error rate is below 0.01 when the resolution is greater than or equal to 8. The outlier with *k* = 2 may be caused by the irregular shape of the 3D models, which seems to have low error rate by accident. In conclusion, the proposed intersection volume computation method achieves high performance in terms of both efficiency and accuracy.

**Figure 9:**
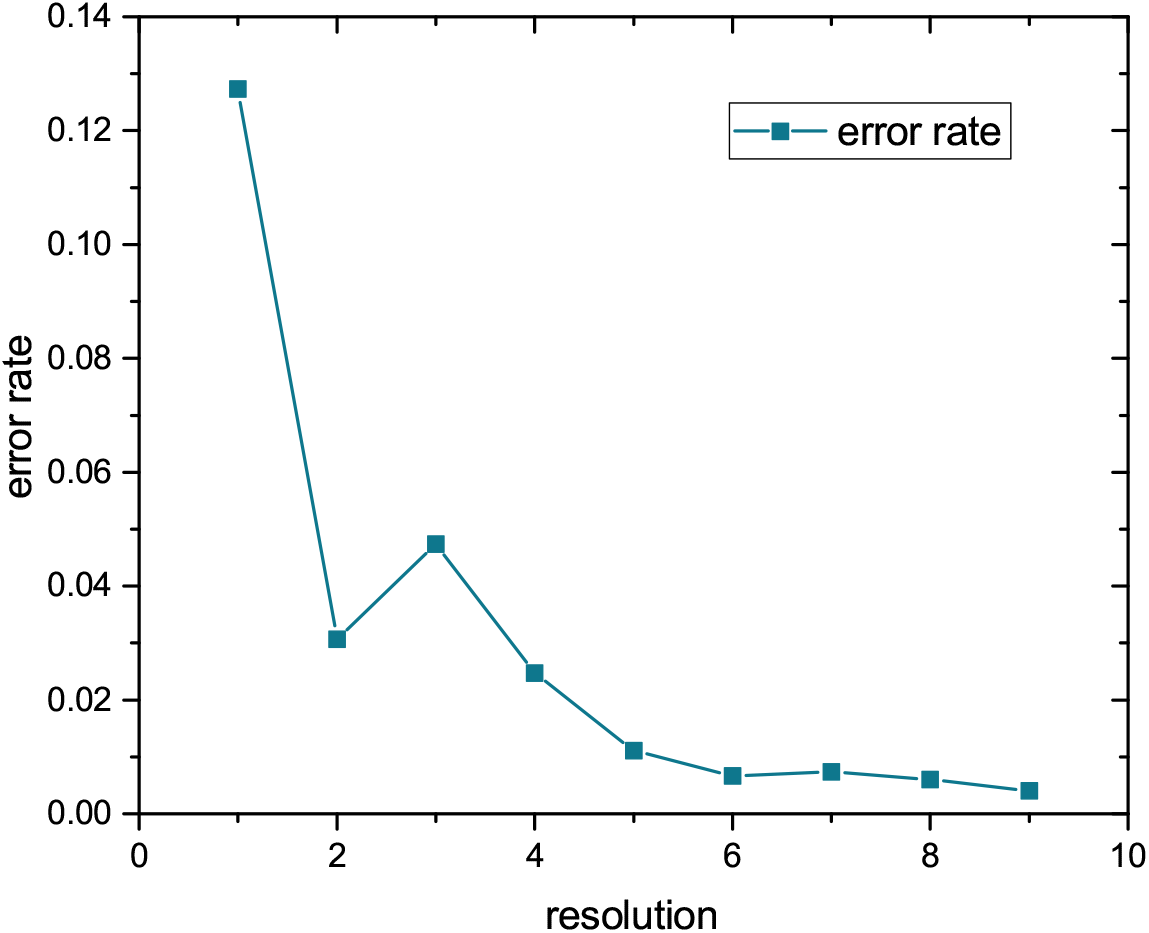
Error rate of different resolutions

## 7 CONCLUSION

We propose a comprehensive framework for 3D tissue block registration for human atlas. In particular, we build multi-level indexing both on MBBs of anatomical structures and primitives of meshes. In addition, we present a voxelization-based method for intersection volume computation. Experimental results show that the framework achieves high accuracy in real-time spatial queries. The 3D registration framework is implemented as a service for HuBMAP, and will be deployed in Amazon AWS platform for production use in short term future.

## 8 Acknowledgments

This research is supported in part by grants from National Institutes of Health 1OT2OD033756-01 and 3OT2OD026671-01S4.

